# Using Bayesian Optimization to Identify Optimal Exoskeleton Parameters Targeting Propulsion Mechanics: A Simulation Study

**DOI:** 10.1101/2021.01.14.426703

**Authors:** GilHwan Kim, Fabrizio Sergi

## Abstract

In this study, we determined the feasibility of modeling the relationship between robot control parameters and propulsion mechanics as a Gaussian process. Specifically, we used data obtained in a previous experiment that used pulses of torque applied at the hip and knee joint, at early and late stance, to establish the relationship a 3D control parameter space and the resulting changes in hip extension and propulsive impulse. We estimated Gaussian models both at the group level and for each subject. Moreover, we used the estimated subject-specific models to simulate virtual human-in-the-loop optimization (HIL) experiments based on Bayesian optimization to establish their convergence under multiple combinations of acquisition functions and seed point selection methods.

Results of the group-level model are in agreement with those obtained with linear mixed effect model, thus establishing the feasibility of Gaussian process modeling. The estimated subject-specific optimal conditions have large between-subject variability in the metric of propulsive impulse, with only 31% of subjects featuring a subject-specific optimal point in the surrounding of the group-level optimal point. Virtual HIL experiments indicate that expected improvement is the most effective acquisition method, while no significant effect of seed point selection method was observed. Our study may have practical effects on the adoption of HIL robot-assisted training methods focused on propulsion.

## I. INTRODUCTION

Robot assisted gait training is becoming a common method for rehabilitation after neurological injury [1]. With the option of mechanical assistance to multiple joints, and several open parameters for the timing of assistance at each joint, controllers for gait training robots are defined by a large number of open parameters, each corresponding to highly variable outcomes of robotic training. To deal with the large number of open parameters in robot-assisted gait training, real-time optimization methods, or Human-In-the-Loop (HIL) optimization methods, have been introduced [2]. In human-robot interaction, HIL methods are used to identify parameters of a robot controller that are optimal in the sense of a specific cost function. In general, this is done by testing the value of the cost function, collecting data to quantify the subject response at those points, and then iteratively updating the controller parameters in real-time to optimize the cost function.

Several successful implementations of HIL optimization exist, many using different optimization algorithms [3]–[4], with one-dimensional (1D) gradient descent method [5], 4D Covariance Matrix Adaptation Evolution Strategy (CMA-ES) [4], and Bayesian optimization [3], [6] used in recent studies using exoskeletons supporting walking function. However, most previous HIL optimization experiments focused on reducing metabolic cost to tune control parameters which results limited applicability in robot-assisted gait training, since the major objective of gait training is to induce desired changes in subjects’ motor coordination with the ultimate goal of improving their motor function. Moreover, on-going research about optimization algorithm is focused on improving performance of Bayesian optimization based on hyperparameter tuning [7], noise modeling [8], acquisition function definition [9], [10], [11], and definition of seed points [3], but it is currently unclear how these methods apply to HIL optimization methods in biomechanics.

In this work, we seek to apply HIL to establish the subject-specific relationship between exoskeleton control parameters and resulting features of gait, such as those describing propulsion mechanics, a crucial component of walking function [12], [13], [14]. We address the feasibility of modeling previously collected data on the effects of the application of pulses of torque to the hip and knee joint on propulsion mechanics using Gaussian process modeling [15]. Specifically, we establish the relationship between pulse torque conditions and propulsion mechanics at the group and subject-specific level, and evaluate how these relationships differ across subjects. Moreover, we run simulations using the estimated subject-specific Gaussian process models to establish convergence in virtual HIL experiments based on Bayesian optimization. Specifically, in our analysis we establish the effect of two important factors, seed point selection method and acquisition function, in the speed and accuracy of convergence of Bayesian optimization targeting propulsion mechanics.

## II. METHODS

### A. Data Collection

In previous research, sixteen healthy subjects participated in an experiment based on the application of pulses of torque to the hip and knee joint to modulate propulsion mechanics [15]. Torque conditions were defined as a combination of three parameters: pulse timing, hip pulse amplitude, and knee pulse amplitude. Two levels were used for pulse timing: pulses were applied at 10% of the estimated gait cycle period (early stance), or 45% of the estimated gait cycle period (late stance). Levels for hip and knee pulse amplitude were defined as either zero torque, flexion, or extension (amplitude was set to 15 N·m for the hip joint, and 10 N·m for the knee joint for both flexion and extension). Sixteen conditions were tested, including all combinations of the factors above, with the exclusion of the combination of zero knee and hip torque. When pulses were applied to both joints, they were applied simultaneously. Each condition was repeated ten times per subject, with a random sequence. Pulses were applied to the right leg during single strides, and spaced by at least eight strides of no pulse application. Hip extension (HE) and propulsive impulse (PI) were assessed at multiple strides: prior to pulse application (stride −1), during pulse application (stride 0), and during the three strides following pulse application (stride 1, 2, 3). In previous work, our group used a linear mixed model to establish the relationship between factors including pulse pattern parameters (pulse timing, amount of torque pulses applied to hip and knee joints), subjects, and stride with propulsion mechanics, as defined by HE and PI [15].

### B. Gaussian Process Modeling

Our first goal is to model the relationship between pulse torque conditions and propulsion mechanics as a Gaussian process. Each subject has 10 measurements of each outcome measure (HE and PI), resulting in a total of 160 data points per stride condition, and thus 800 data points per subject (12800 total measurements), referred to as measurements *y*_*HE*_ and *y*_*P I*_. These measurements can be indexed as a function of factors: pulse timing (*T*), amplitude of hip and knee torque pulses (*K* and *H*), stride (*S*), subject (*Sbj*), trial index (*Rep*), as

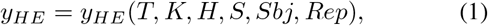

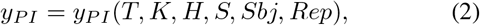

where *T* ∈ (10, 45), *K* ∈ (−10, 0, 10), *H* ∈ (−15, 0, 15), *S* ∈ (−1, …, 3), *Sbj* ∈ (1, …, 16), and *Rep* ∈ (1, …, 10).

We assumed that the Gaussian process model linking control parameters with the outcome measures of propulsion should follow the characteristics listed below:

- noise from outcome measures is normally distributed with zero mean 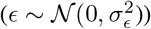;
- noise is independent of the human response;
- the distribution of the human response under repeated exposure to the same pulse condition is normal;
- the variance of the human response will be constant under different pulse conditions.

#### 1) Group-level Model Formulation

Data in variables (1) and (2) are concatenated along the subject dimension obtaining variables 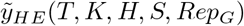 for HE, and 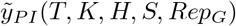 for PI, with 160 repetitions available for factor *Rep*_*G*_.

The modified outcome measures (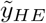 and 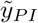) can be expressed as the sum of an unknown Gaussian process *G*, and noise (*ϵ*) as follows:

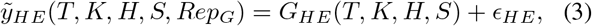

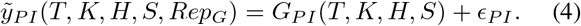

The average in the measurements 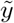 can be derived by average value among 160 repetitions per each condition. Since noise is assumed to have zero mean, the average value of the unknown Gaussian process *G* is equal to the average in the measurements 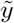. The variance in the measurements 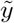 are expressed as a sum of model variance (variations in the true output arising from repeated exposure to the same conditions 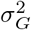), and measurement error (variations in the measurements that are not associated to changes in the true value of the output 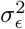). Since the noise is constant for all values of pulse torque factors, the following relationships hold true:

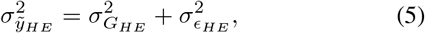

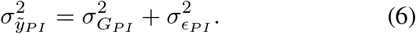

These relationships are used to specify values for the model variance 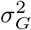 used for the Gaussian process covariance function. Values of 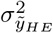 and 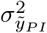 are calculated as the maximum variance of 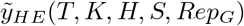 along the *Rep*_*G*_ dimension, i.e. the one resulting from the combination of pulse parameters associated with the largest variance across subjects and repetitions.

#### 2) Group-level Model Estimation

A Gaussian process model is estimated from eq. (3) and (4) to approximate the mean and variance of data 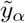. In a Gaussian process, the variance is defined based on a covariance (or kernel) function *k*_*α*_(*x*_*i*_, *x*_*j*_), which defines how variance propagates within a dimension, or in different dimensions of the model. In this work, we defined the covariance function (automatic relevance determination squared exponential kernel) as:

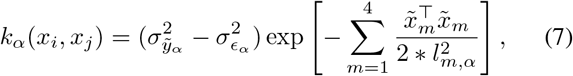

where *x* is set of pulse torque input parameters, *x* = (*T, K, H, S*), 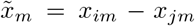 *l*_*m,α*_ is the *m*-th length scale hyperparameter, *α* ∈ (*HE, PI*). (*i, j*) are indices corresponding to two arbitrary points in the 4D domain of the parameters. Using the covariance function (7), the estimated mean value and variance of a point *x*_*n*+1_, based on *n* measurements, are

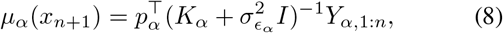

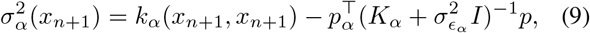

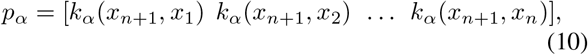

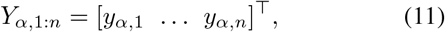

and

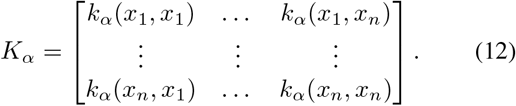

In our case, *n* = 80, referring to the fact that all observations (*Y*_*α,*1:*n*_) are used to estimate a Gaussian process model for *G*_*HE*_ and *G*_*P I*_. After a model is estimated given the available *n* measurements, the estimated average value *μ*(*x*_*n*+1_) of all points in the domain is defined as the mean function *m*(*x*) of input *x*.

As discussed above, (3) and (4) define the relationship between control parameters and output using Gaussian processes, i.e., 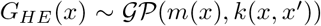, where *k*(*x, x*′) is the kernel function. As such, the Gaussian processes can be estimated by solving the following least-squares problems in terms of the hyperparameters *l*_*m,α*_ and 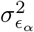:

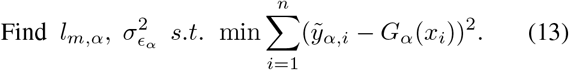

To align our fitted model with observations emerging from our previous linear mixed model analysis, we set constraints on the hyperparameters optimized in (13) (Table I).

**TABLE 1.** LOWER AND UPPER BOUNDS FOR ESTIMATED NOISE STANDARD DEVIATION AND FOUR LENGTH SCALE HYPERPARAMETERS

| | T [% gait cycle] | S [] | K [N.m] | H [N.m] | $\sigma_{\epsilon_\alpha}$ |
| --- | --- | --- | --- | --- | --- |
| LB | 12 | 0 | 6 | 7 | 0 |
| UB | 40 | 1 | 15 | 20 | $\max(\sigma_{\tilde{y}_\alpha})$ |

#### 3) Subject-specific Model Formulation

Similarly as the group data, the outcome measures for each subject (*y*_*Sbj,HE*_ and *y*_*Sbj,P I*_) can be expressed as the sum of an unknown subject-specific Gaussian process *G*_*Sbj*_, and noise (ϵ_*Sbj*_):

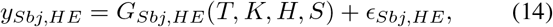

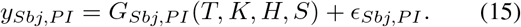

Measurements collected from each subject correspond to the ten repeated measurements, collected at all combinations of the factors, resulting in 800 measurements. Similarly as done in the group-level model, noise is assumed to be constant for all values of pulse torque factors, hence variances of data and processes are linked by:

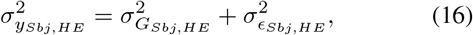

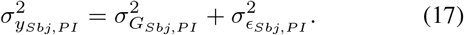

#### 4) Subject-specific Model Estimation

Similarly as done for group-level model estimation, a Gaussian process is estimated from (14) and (15). The same kernel function defined in (7) was used for the definition of the subject-specific Gaussian models, with the difference that 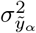 was defined as the variance resulting from the combination of pulse parameters with the largest within-subject variance across the ten repetitions. Because it is usually impractical to collect a sufficient number of observations from an individual subject to properly estimate subject-specific length-scale hyperparameters, we proceeded to specify for individuals the same values of parameters *l*_*m,α*_ estimated for the group. In these conditions, the subject-specific Gaussian processes *G*_*Sbj,α*_ are estimated solving the following least-squares problem:

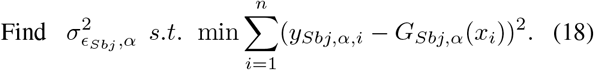

#### 5) Quantifying Variability of Maxima in Subject-specific Models

To quantify the variability of the optimal points in the estimated subject-specific Gaussian process models *G*_*Sbj,HE*_ and *G*_*Sbj,P I*_, we calculated the vectors 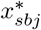 as the values of *T*, *K*, and *H* that maximized the estimated process value at stride 0 for different subjects, and 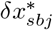 as the values of *T*, *K*, an *H* that maximized the estimated change in outcome measure between stride 0 and stride −1 for different subjects. We thus established for how many subjects (*n*_*w*_) the subject-specific optimal points fell within a sphere of radius *r* centered around the optimal points estimated using the group-level model (*x*\* and *δx*\*). The optimal point analysis is conducted on the non-dimensional domain where all coordinates are comprised between 0 and 1 using a min-max normalization. The functions *n*_*w*_(*r*) are compared to establish variability of maxima in the two outcome measures.

### C. Bayesian Optimization Simulation

Virtual HIL experiments are conducted to determine the convergence of Bayesian optimization for each subject-specific Gaussian process model, under different settings of the optimization algorithm. Each subject-specific Gaussian process model is assumed to be the human response (i.e., data generating process) when pulse torque assistance is applied. Three robotic control parameters (*T*, *K*, and *H*) are used as input parameters. In all conditions, the optimization algorithm does not have any knowledge about the subject-specific model at the beginning of each optimization. Two algorithm components are tested using a factorial design: i) acquisition function (three levels), and *ii)* and seed point selection method (three levels) are evaluated in terms of the number of iterations required for convergence. In total, the 9 combinations of the two components are tested in simulation, with twenty simulations repeated for each combination for each subject.

#### 1) Acquisition Function

Expected Improvement (EI), Probability of Improvement (PoI), and Lower Confidence Bound (LCB) are used in this study. EI finds the next point which maximizes mean value improvement. This expected improvement is defined as

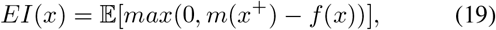

where *x*^+^ is the best point from data points explored so far and *m*(*x*^+^) is mean value of Gaussian process model value at point *x*^+^. PoI finds the next point which has the maximum probability of improvement as

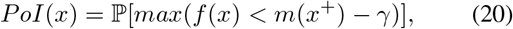

where *γ* is margin. The LCB acquisition function selects the next point which maximizes the lower bound as

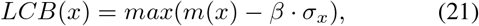

where *σ*_*x*_ is standard deviation at point *x*, and *β* > 0 is a heuristic trade-off (*β* = 2 in this work).

#### 2) Seed Point Selection Method

Three seed point selection methods are used in this work. In the first method (*divide*), eight points are selected by dividing the parameter search space in eight regions (the domain for each parameter is divided into two equal parts), and then each of the eight points are selected randomly within each of the eight regions. In the second method (*random*), eight points are randomly defined in the search space. In the third method (*optimal*), the set of eight seed points is composed of seven random points in the search space, plus the optimal point of the group-level Gaussian process model.

#### 3) Virtual Optimization

Virtual Bayesian optimization is conducted to maximize HE and PI at the stride of pulse application, and the outcome measure is defined as the number of iterations *n*_*x*_ and *n*_*y*_ required to achieve one of two convergence criteria.

If 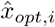 is the normalized coordinates optimal point estimated via virtual optimization after *i* iterations, and *x*_*opt*_ is the known optimal point for that subject, *n*_*x*_ is defined as the minimum value of *i* where the normalized difference *x*_*diff*_ between the two quantities is smaller than 10%, where

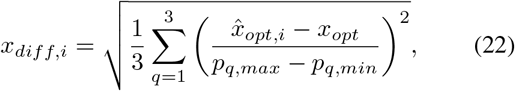

where *p* is the range of each component of *x*.

The second criterion is based on the value of the estimated outcome after a certain number of iterations. Specifically, *n*_*y*_ is defined as the minimum number of iterations where the normalized estimation error *f*_*diff*_ (*x*_*i*_) is smaller than 5% of the known subject-specific model maximum *G*_*sbj,α*_(*x*\*), where

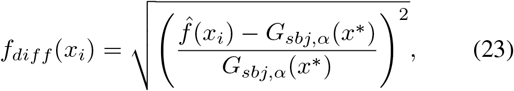

All simulations were run for 80 iterations. For each criterion, if no convergence was achieved within 80 simulations, *n* was set to 80.

Four two-way ANOVA, one for each outcome measure (combination of convergence criterion – *n*(*x*) and *n*(*y*)) – and propulsive metric – HE and PI – were conducted to quantify the effects of the two factors (i.e., acquisition function, three levels, and seed point selection methods, three levels) on convergence speed.

## III. RESULTS

### A. Gaussian Process Modeling

#### 1) Group-level Model

Results for group-level Gaussian process modeling are shown in Fig. 1 and Fig. 2. The process variances 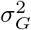 calculated from group-level propulsion are equal to 4.31 deg2 for HE and 2.12 N2s2. Estimated noise standard deviation and hyperparameter values are listed in Table II.

**Fig. 1.**
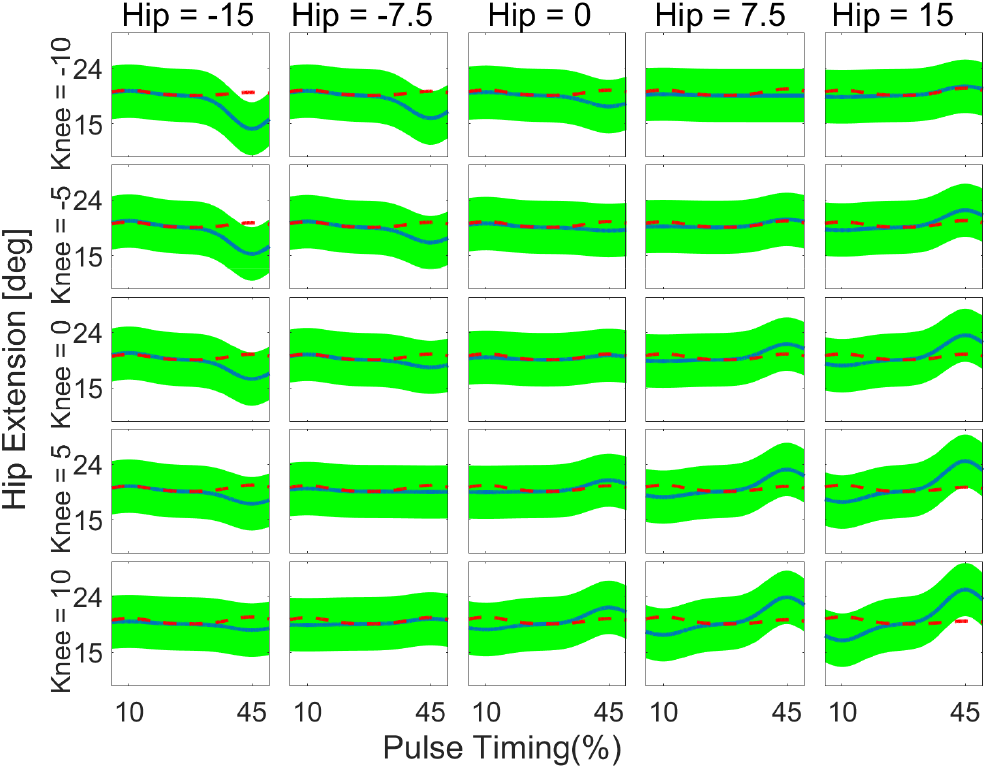
Gaussian process model of group-level Hip Extension (HE) data. Pulse factors are indexed by the horizontal panel coordinate (hip torque), vertical panel coordinate (knee torque), *x* axis within a panel (pulse timing). The two lines in each panel indicate different levels of factor stride. Red dashed line: model at the stride before intervention (stride 1); blue line: model at the stride of intervention. Hip and Knee indicate the amplitude of torque pulses applied to hip joint and knee joint respectively. Extension torques are positive in hip and knee. At each point, shaded areas extend one standard deviation beyond the mean.

**Fig. 2.**
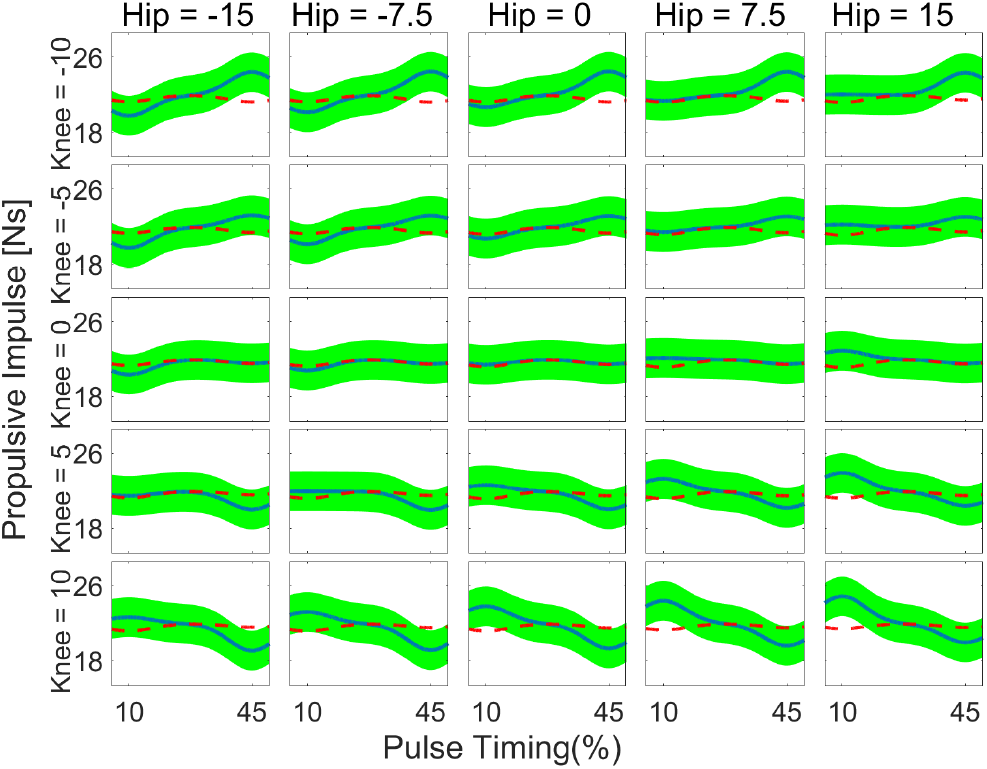
Gaussian process model of group-level Propulsive Impulse (PI) data.

**TABLE 2.** LENGTH SCALE HYPERPARAMETER VALUES AND ESTIMATED NOISE STANDARD DEVIATION

| | T [% gait cycle] | S [] | K [N·m] | H [N·m] | $\sigma_{\epsilon_\alpha}$ |
| --- | --- | --- | --- | --- | --- |
| HE | 26.71 | 0.73 | 6.17 | 7.43 | 0.21 deg |
| PI | 12.18 | 0.50 | 8.66 | 11.73 | 0.78 Ns |

#### 2) Subject-specific Results

The mean of the distribution of subject-specific process variances 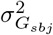 were equal to 69.04 deg^2^ and 33.86 N^2^s^2^ for HE and PI, respectively. The number of optimal points for subject-specific models within percentage difference of the group-level model is shown in Fig. 5. The optimal points are for maximal outcomes during stride 0 and maximum change in outcome (between stride 0 and stride −1) for each subject-specific model. The optimal points for the subject-specific models are shown in Fig. 3 and Fig. 4 for HE and PI, respectively. 12 subject-specific Gaussian process models have optimal points that have a percentage difference of less than 20 % compared to the optimal point of group-level Gaussian process model for HE (Fig. 3). For propulsive impulse case (Fig. 4), only 5 optimal points of subject-specific Gaussian process model are located within 20% difference from the optimal point of group-level Gaussian process model. For PI, the minority of subjects exhibited subject-specific optimal points within a reasonable neighborhood of the group-level optimal point. In fact, more optimal points of subject-specific models in PI case are located in late stance (45 %) than early stance (10 %) where optimal point of group-level model is located.

**Fig. 3.**
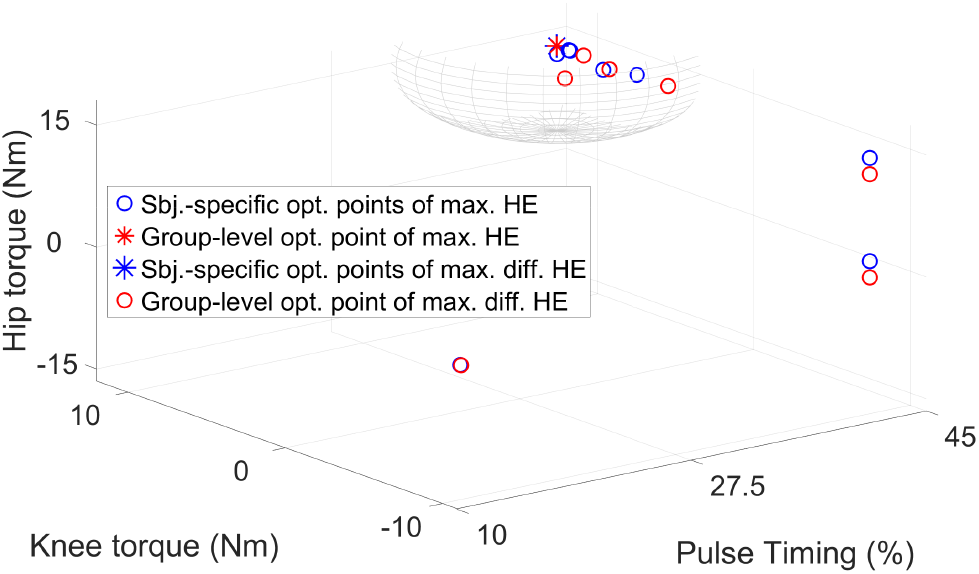
Distribution of the optimal points of subject-specific models (circles) and of the group-level model (asterisks) for HE. The 20% range is indicated by a sphere centered around the optimal point of the group-level model.

**Fig. 4.**
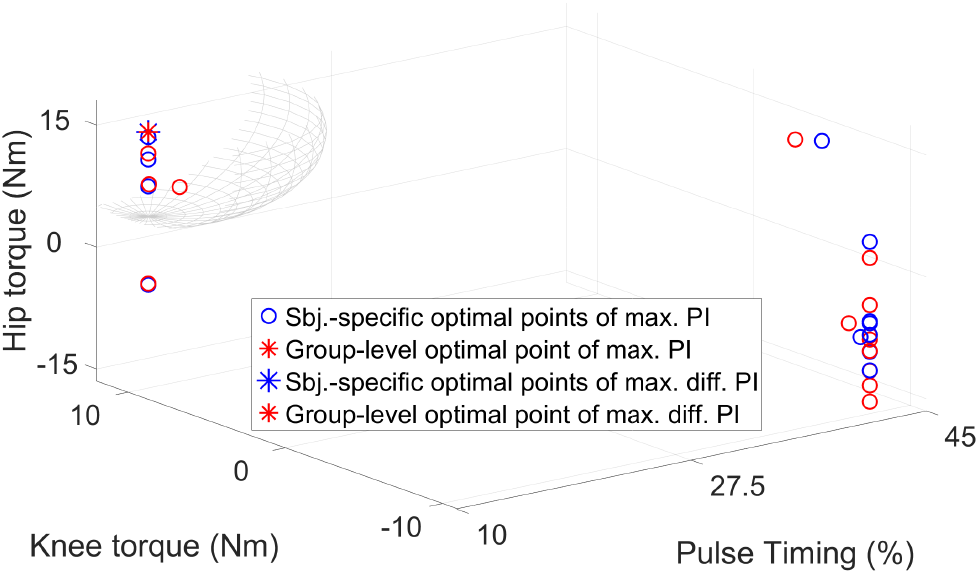
Distribution of the points of sixteen subject-specific models (circles) and of the group-level model (asterisk dots) for PI. 20 % difference are indicated by grid sphere around optimal point of group-level model.

**Fig. 5.**
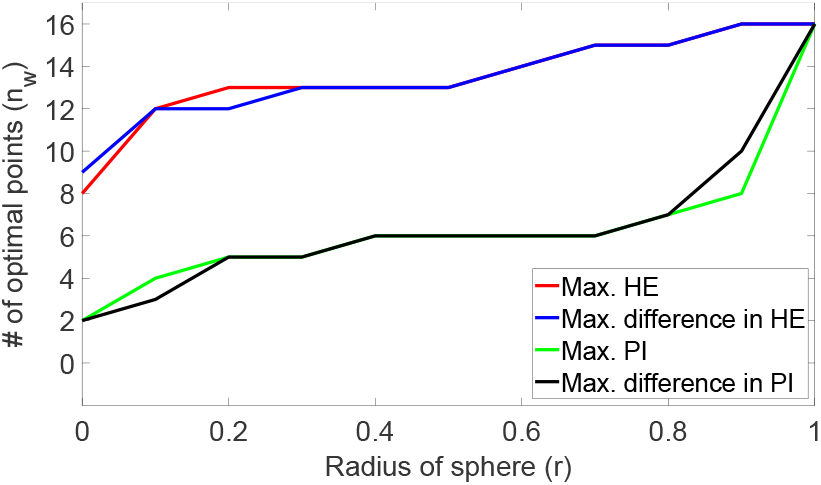
Number of subjects (*nw*) whose subject-specific model maxima are within a specified distance (*r*) from the group-level model maximum.

### B. Bayesian Optimization Simulation

Virtual Bayesian optimization experiments achieved convergence within 80 iterations in 80.58 % of runs (94.40 ± 6.06 % when using EI, 85.57 ± 13.94 % when using PoI, 61.77 ± 26.66 % when using LCB). The results of the ANOVA analysis are reported in Table III and Fig. 6. Based on the ANOVA analysis, the seed point selection method had no significant effect on the outcome measure in any condition. Instead, the choice of an acquisition function showed to be associated with the number of iterations required for convergence (*p* < 0.001 in all conditions tested). Specifically, EI was the only acquisition function type that was estimated to have negative coefficient in the linear model when using *random* method and PoI as neutral level of the factors, indicating that its effect is to decrease the number of iterations to obtain convergence. Post-hoc tests are conducted to distinguish the significant difference among groups under different combinations of acquisition functions and seed point selection methods (Fig. 6). Post-hoc tests indicate that within the same group of acquisition function types, there exist no significant differences (*p* > 0.001 for all comparisons). Moreover, post-hoc tests comparing different acquisition functions showed that EI was significantly better than LCB in all conditions tested, and afforded a significantly greater accuracy in identifying optimal control parameters compared to PoI.

**TABLE 3.** TWO WAY ANOVA TEST RESULTS FOR NUMBER OF ITERATIONS TO ESTABLISH CONVERGENCE

| Source | HE |  |  | Point location (HE) |  |
| --- | --- | --- | --- | --- | --- |
|  | F ratio | Prob> \|F\| |  | F ratio | Prob> \|F\| |
| Acq. func. | 27.9950 | < 0.001 |  | 150.1179 | < 0.001 |
| Seed pt. | 0.0702 | 0.9322 |  | 0.2270 | 0.7969 |
| Interaction | 0.2350 | 0.9187 |  | 1.2166 | 0.3017 |

| Source | PI |  |  | Point location (PI) |  |
| --- | --- | --- | --- | --- | --- |
|  | F ratio | Prob> \|F\| |  | F ratio | Prob> \|F\| |
| Acq. func. | 100.9153 | < 0.001 |  | 137.0745 | < 0.001 |
| Seed pt. | 0.2063 | 0.8136 |  | 0.1297 | 0.8784 |
| Interaction | 0.1021 | 0.9818 |  | 0.5217 | 0.7198 |

**Fig. 6.**
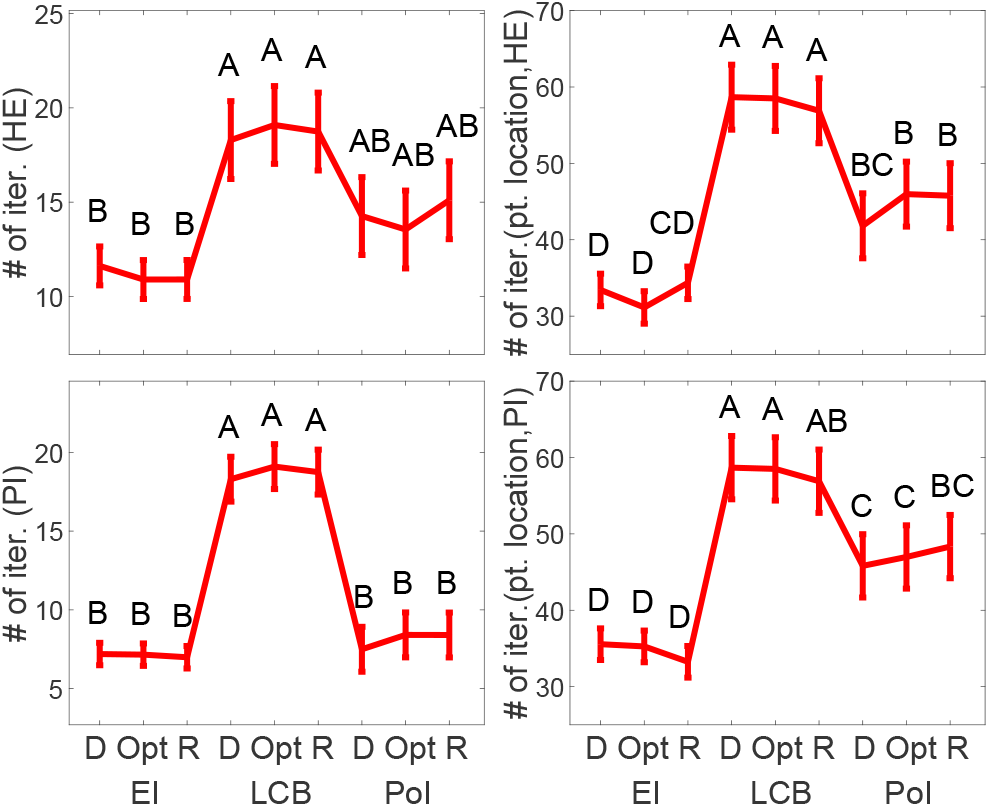
2-way ANOVA interaction plot with post-hoc test results. D, Opt, and R in x-axis indicate the *divide* method, *optimal* method, and *random* method for seed point selection method respectively. Pairs of groups that share the same letter label are not significantly different.

## IV. DISCUSSION AND CONCLUSIONS

In this study, we determined the feasibility of using Gaussian process modeling to establish modeling the relationship between three exoskeleton control parameters (hip torque pulse amplitude, knee torque pulse amplitude, and timing of the pulses), and outcome measures relevant for propulsion. To estimate parameters of a Gaussian process, we estimated the noise variance and tuned the hyperparameters of a Gaussian process describing the response at the group level. Based on the group-level Gaussian process model, we estimated 16 subject-specific models and used those as data-generating process for virtual HIL Bayesian optimization simulations. In the virtual Bayesian optimizations, we established how many iterations would be required for convergence of a hypothetical experiment, and quantified the effects of different acquisition functions and seed point selection methods on convergence speed.

Our results of group-level Gaussian process model (Fig. 1 and Fig. 2) demonstrate that it is feasible to construct a model between the 3D space of control parameters and outcomes using a Gaussian process with estimated noise variance. Based on previous predictions using linear mixed model [15], group-level Gaussian process models for both HE and PI are expected to have a similar response when pulses are applied to the knee and hip joints. In agreement with the results of the previous study [15], hip extension torque increases HE in late stance but decreases HE in early stance; knee extension and hip extension pulse torque increase PI in early stance; knee extension pulse torque decreases PI in late stance. The variability of optimal points for subject-specific models is moderate in HE and high in PI. Specifically for the PI case, the maxima of subject-specific models fell within 20% normalized distance from the optimal point for group-level model only in the minority of cases (5/16). This indicates that the optimal point for group-level model is likely to be distant from the optimal point for subject-specific models. In this case, in fact, optimal points for subject-specific models are separated to make two clusters of optimal points in early and late stance. Our analysis in HE case is aligned with previous work targeting reduction in metabolic cost, indicating that one subject’s optimal point can be optimal for another subject [3].

In contrast with previous research selecting seed points by dividing the search space in *N* regions and randomly selecting seed points from all the *N* regions [3], seed point selection method is not estimated to make a significant difference in a single pulse torque application experiment (*p* > 0.12 for the different conditions tested, Table III). This results are consistent in both HE and PI cases. Since we included *optimal* method in seed point selection method, this results also indicate that optimal point location relationship between group-level optimal point and subject-specific optimal points in HE case do not effect results of simulations. However, the type of acquisition function used for optimization did significantly contribute to the number of iterations required to achieve convergence in all cases of outcome measures (HE and PI) and point locations (*p* < 0.001). Since the interaction term between types of acquisition function and seed point selection methods is not significant (*p* > 0.1 for all conditions tested), the optimal acquisition function (in terms of convergence speed) of HIL Bayesian optimization is estimated to be EI. Therefore, our expectation toward HIL Bayesian optimization setting is that LCB will require the largest number of iterations to establish convergence, while EI will require the smallest number for iterations for convergence regardless of seed point selection methods.

This study has limitations. Since virtual HIL Bayesian optimization rely on our assumption that subject-specific Gaussian process models can fully represent actual human response, the validity of these models directly relates to the validity of the results of Bayesian optimization obtained in this study. Another limitation is that noise is not fully eliminated to construct a model that relate control input parameters and propulsive mechanics. As shown in Fig. 1 and Fig. 2, the model at the stride before intervention (red dashed line) have changes with control input parameters, which is impossible as a change in output cannot anticipate a change in input. This result indicates that our optimization procedure for estimating hyperparameters and noise variance may need to be improved, or that the relationship between input parameters and improvement of outcome measures (HE and PI) should be considered to cancel out noise from stride at intervention (stride 0) and stride before intervention (stride −1) Gaussian process models.

Overall, our study identified a new method based on Gaussian process modeling to estimates the relationship between exoskeleton control parameters and specific gait features (HE and PI), and established the effects of seed point selection method and acquisition function types in number of iterations for convergence. These results make foundation of promising approach for planning further HIL robotic gait training experiment by focusing on specific gait features to target with training.

